# The *Drosophila* RNA binding protein Nab2 patterns dendritic arbors and axons via the planar cell polarity pathway

**DOI:** 10.1101/2021.11.10.468151

**Authors:** Edwin B. Corgiat, Sara M. List, J. Christopher Rounds, Dehong Yu, Ping Chen, Anita H. Corbett, Kenneth H. Moberg

## Abstract

RNA binding proteins support neurodevelopment by modulating numerous steps in post-transcriptional regulation, including splicing, export, translation, and turnover of mRNAs that can traffic into axons and dendrites. One such RBP is ZC3H14, which is lost in an inherited intellectual disability. The *Drosophila melanogaster* ZC3H14 ortholog, Nab2, localizes to neuronal nuclei and cytoplasmic ribonucleoprotein granules, and is required for olfactory memory and proper axon projection into brain mushroom bodies. Nab2 can act as a translational repressor in conjunction with the Fragile-X mental retardation protein homolog Fmr1 and shares target RNAs with the Fmr1-interacting RBP Ataxin-2. However, neuronal signaling pathways regulated by Nab2 and their potential roles outside of mushroom body axons remain undefined. Here, we demonstrate that Nab2 restricts branching and projection of larval sensory dendrites via the planar cell polarity pathway, and that this link may provide a conserved mechanism through which Nab2/ZC3H14 modulates projection of both axons and dendrites. Planar cell polarity proteins are enriched in a Nab2-regulated brain proteomic dataset. Complementary genetic data indicate that Nab2 guides dendrite and axon growth through the planar-cell-polarity pathway. Analysis of the core planar cell polarity protein Vang, which is depleted in the *Nab2* mutant whole-brain proteome, uncovers selective and dramatic loss of Vang within axon/dendrite-enriched brain neuropil relative to brain regions containing cell bodies. Collectively, these data demonstrate that Nab2 regulates dendritic arbors and axon projection by a planar-cell-polarity-linked mechanism and identify Nab2 as required for accumulation of the core planar cell polarity factor Vang in distal neuronal projections.

## Introduction

While many key developmental events are triggered by extracellular factors that signal through cytoplasmic cascades to alter nuclear gene transcription, other key events are triggered by shifts in posttranscriptional processing or localization of mRNAs that guide cell fates and differentiation. Importantly, the fidelity of these mRNA-based developmental mechanisms relies on RNA binding proteins (RBPs) that associate with nascent RNAs and regulate splicing, export, stability, localization, and translation (Schieweck *et al*. 2021). These key regulatory mechanisms are particularly evident in the developing nervous system, where mutations in genes encoding RBPs are often linked to human diseases. Examples of this linkage include Fragile X mental retardation protein (FMRP) (Gross *et al*. 2012), the survival of motor neuron protein (SMN) (Edens *et al*. 2015), and the TAR DNA binding protein 43 (TDP-43) (Agrawal *et al*. 2019; Gebauer *et al*. 2021). Sensitivity of the central and peripheral nervous systems to loss of RBPs has been attributed to the importance of post-transcriptional mechanisms, such as local translation of mRNAs and brain-specific extension of 3’UTRs (Mattioli *et al*. 2017; Thelen and Kye 2019; Engel *et al*. 2020) that enable fine-tuned spatiotemporal control of neuronal gene expression. This spatiotemporal control of mRNA processing and translation plays an important role in forming complex dendritic architectures and the uniquely polarized morphology of neurons (Lee *et al*. 2003). Accordingly, neurological diseases caused by mutations in genes encoding RBPs often include defects in axonal or dendritic morphology (Jung *et al*. 2012; Hornberg and Holt 2013; Holt *et al*. 2019), and in some cases, these axonal and dendritic defects can be traced to defective post-transcriptional control of one or a few mRNAs normally bound by the corresponding RBP.

The human *ZC3H14* gene encodes a ubiquitously expressed zinc-finger, polyadenosine RBP (<u>Z</u>nF <u>C</u>ys<u>C</u>ys<u>C</u>ys<u>H</u>is #14) that is lost in an inherited form of intellectual disability (Pak *et al*. 2011). Studies in multiple model organisms have begun to define functions for ZC3H14 in guiding neuronal morphogenesis. Analysis of the sole *Drosophila* ZC3H14 homolog, Nab2, detects cell-autonomous requirements in Kenyon cells (KCs) for olfactory memory as well as axonal branching and projection into the brain mushroom bodies (MBs) (Kelly *et al*. 2016; Bienkowski *et al*. 2017), twin neuropil structures that are the center for associative olfactory learning in insects (Thum and Gerber 2019). Significantly, transgenic expression of human ZC3H14 only in fly neurons is sufficient to rescue a variety of *Nab2* null phenotypes (Pak *et al*. 2011; Kelly *et al*. 2014; Kelly *et al*. 2016), supporting a model in which Nab2 and ZC3H14 share critical molecular roles and mRNA targets. The *Zc3h14* gene is not essential in mice but its loss results in defects in working memory (Rha *et al*. 2017) and dendritic spine morphology (Jones *et al*. 2021). An accompanying proteomic analysis of *Zc3h14* knockout hippocampi identified several proteins involved in synaptic development and function that change in abundance upon ZC3H14 loss (Rha *et al*. 2017), and which are thus candidates to contribute to *Zc3h14* mutant phenotypes. Intriguingly, the homologs of some of these ZC3H14-regulated proteins in the mouse hippocampus are also sensitive to Nab2 loss in the developing *Drosophila* pupal brain (Corgiat *et al*. 2021), suggesting conserved links between Nab2 and ZC3H14 and neurodevelopmental pathways.

A variety of intercellular signaling mechanisms play required roles in sensing extracellular cues that guide the complex axonal and dendritic structures that characterize specific areas of the central and peripheral nervous system (CNS and PNS). These cascades can respond to long-range directional cues, such as Netrin signaling, or to short-range directional cues from the Slit-Robo, Abl-Ena, and Semaphorin pathways (Puram and Bonni 2013; Stoeckli 2018). One pathway with an emerging role in both axonal and dendritic development is the planar cell polarity (PCP)-noncanonical Wnt pathway (Zou 2004; Andre *et al*. 2012; Zou 2012; Gombos *et al*. 2015; Misra *et al*. 2016). PCP signals are based on asymmetric distribution of two apically localized transmembrane complexes, which in *Drosophila* correspond to the Stan-Vang-Pk complex (Starry Night aka Flamingo-Van Gogh-Prickle) and the Stan-Fz-Dsh-Dgo complex (Frizzled-Disheveled-Diego); these complexes are intracellularly antagonistic but intercellularly attractive, leading to apical polarization across an epithelial plane (Taylor *et al*. 1998; Boutros and Mlodzik 1999; Vladar *et al*. 2009; Goodrich and Strutt 2011;Adler 2012; Peng and Axelrod 2012; Adler and Wallingford 2017; Mlodzik 2020). Core PCP components signal to downstream effector molecules that exert localized effects on the F-actin cytoskeleton (Courbard *et al*. 2009; Adler 2012; Soldano *et al*. 2013; Fagan *et al*. 2014; Gombos *et al*. 2015), which in turn guides epithelial traits like proximal-distal wing hair orientation in *Drosophila* and sensory hair cell polarity in the mouse cochlea (Jones and Chen 2007; Qian *et al*. 2007; Simons and Mlodzik 2008; Chacon-Heszele and Chen 2009; Rida and Chen 2009; Alpatov *et al*. 2014). One such factor is encoded by the *β amyloid protein precursor-like* (*Appl*) gene and modulates the PCP pathway during axonal and dendritic outgrowth (Soldano *et al*. 2013; Liu *et al*. 2021). Importantly, PCP is required for axon guidance in specific groups of neurons in *Drosophila, C. elegans*, mice, and chick, and for dendritic branching of mouse cortical and hippocampal neurons, and *Drosophila* body wall sensory neurons (Hindges *et al*. 2002; Mclaughlin and O’leary 2005; Schmitt *et al*. 2006; Shafer *et al*. 2011; Cang and Feldheim 2013; Yoshioka *et al*. 2013; Hagiwara *et al*. 2014; Yasumura *et al*. 2021). For example, loss of the murine *Vang* homolog *Vangl2* leads to defects in axon guidance of spinal cord commissural axons (Shafer *et al*. 2011), and *dsh* mutants in *C. elegans* cause neuronal projection and morphology defects (Zheng *et al*. 2015). In *Drosophila*, loss of the core PCP components *stan, Vang, pk, fz*, or *dsh* individually disrupt α and β axon projection into the MBs (Shimizu *et al*. 2011; Ng 2012). Intriguingly, loss of *stan* or its LIM-domain adaptor *espinas* (*esn*) also disrupts dendritic self-avoidance among the class IV dendritic arborization (da) neurons (Matsubara *et al*. 2011), demonstrating a requirement for PCP factors in both axon and dendrite morphogenesis within sets of neurons in the central (CNS) and peripheral (PNS) nervous systems.

Integrating data from two of our recent studies provides evidence for pathways through which the Nab2 RBP could guide axonal and dendritic projections. These analyses, one a genetic modifier screen based on a *GMR-Nab2* rough eye phenotype (Lee *et al*. 2020) and the other a proteomic analysis of *Nab2* null pupal brains (Corgiat *et al*. 2021), each suggest a link between Nab2 and the PCP pathway. The *GMR-Nab2* modifier screen identified alleles of PCP components, both core components and downstream effectors (e.g., *Vang, dsh, fz, stan, pk, Appl*, and the formin *DAAM*), as dominant modifiers of *Nab2* overexpression phenotypes in the retinal field (Lee *et al*. 2020). In parallel, gene ontology (GO) analysis of proteomic changes in *Nab2* null brains detected enrichment for dendrite guidance and axodendritic transport GO terms among affected proteins (Corgiat *et al*. 2021), which include the core PCP factor Vang and the PCP accessory factor A-kinase anchor protein 200 (Akap200). Significantly, *Drosophila* Vang and its murine homolog Vangl2 are one of six pairs of homologs whose levels change significantly in *Nab2* null fly brains and *Zc3h14* knockout mouse hippocampi (Rha *et al*. 2017; Corgiat *et al*. 2021), suggesting a conserved relationship between Nab2/ZC3H14 and the PCP pathway in the metazoan central nervous system (CNS).

Considering observations outlined above, we have investigated interactions between Nab2 and PCP factors in two neuronal contexts - CNS axons of the *Drosophila* pupal MB α- and β-lobes, and in larval dendrites of class IV dorsal dendritic arbor C (ddaC) neurons - which provide complementary settings to analyze the Nab2-PCP link in axonal and dendritic compartments. We detect enrichment for PCP factors among brain-enriched proteins affected by Nab2 loss and a pattern of genetic interactions between *Nab2* and multiple PCP alleles in both MB axons and ddaC dendrites that are consistent with Nab2 regulating axon and dendrite outgrowth by a common PCP-linked mechanism. However, differences in how individual PCP alleles modify axonal vs dendritic *Nab2* mutant phenotypes suggest that the Nab2-PCP relationship may vary between neuronal subtypes (i.e., pupal Kenyon cells vs. larval ddaC neurons). Cell type-specific RNAi indicates that Nab2 acts cell autonomously to guide PCP-dependent axon and dendrite growth, implying a potentially direct link between Nab2 and one or more PCP components within Kenyon cells and ddaC neurons. Based on the drop in Vang levels detected by proteomic analysis of *Nab2* null brains (Corgiat *et al*. 2021), we analyze the levels and distribution of a fluorescently tagged Vang protein in adult fly brains. Consistent with prior bulk proteomic data, overall Vang-GFP fluorescence is reduced in *Nab2* null brains compared to control; significantly, this drop is accompanied by an unexpected and selective loss of Vang protein in neuropil regions enriched in axons compared to regions enriched in cell bodies. Collectively, these data demonstrate that Nab2 is required to regulate axonal and dendritic growth through a PCP-linked mechanism and identify the Nab2 RBP as required for the accumulation of Vang protein into distal axonal compartments.

## Materials and Methods

### *Drosophila* genetics

All crosses were maintained in humidified incubators at 25°C with 12hr light-dark cycles unless otherwise noted. The *Nab2*^*ex3*^ loss of function mutant has been described previously (Pak *et al*. 2011). Alleles and transgenes: *Nab2*^*EP3716*^ (referred to as “*Nab2 oe*”; Bloomington (BL) #17159), *UAS-Nab2*^*RNAi*^ (Vienna *Drosophila* Research Center (VDRC), #27487), *UAS-fz2*^*RNAi*^ (BL #27568), *appl*^*d*^ (BL #43632), *dsh*^*1*^ (BL #5298), *Vang*^*stbm-6*^ (BL #6918), *pk*^*pk-sple-13*^ (BL #41790), *Vang*^*EGFP*.*C*^ (‘*Vang-eGFP’)* (gift of D. Strutt), *ppk-Gal4;UAS-mCD8::GFP* (gift of D. Cox), and *w*^*1118*^ (‘control’).

### *Drosophila* brain dissection, immunohistochemistry, visualization, and statistical analysis

Brain dissections were performed essentially as previously described (Kelly *et al*. 2016). Briefly, 48-72 hours after puparium formation (APF) brains were dissected in PBS (1xPBS) at 4°C, fixed in 4% paraformaldehyde at RT, washed 3x in PBS, and then permeabilized in 0.3% PBS-T (1xPBS, 0.3% TritonX-100). Following blocking for 1hr (0.1% PBS-T, 5% normal goat serum), brains were stained overnight in block+primary antibodies. After 5x washes in PBS-T (1xPBS, 0.3% TritonX-100), brains were incubated in block for 1hr, moved into block+secondary antibody for 3hrs, then washed 5x in PBS-T and mounted in Vectashield (Vector Labs). Antibodies used: anti-FasII 1D4 (Developmental Studies Hybridoma Bank) at 1:50 dilution, anti-GFP polyclonal (ThermoFisher Catalog# A-11122) at a 1:200 dilution, and anti-nc82 (Developmental Studies Hybridoma Bank) at 1:50 dilution. Whole brain images were captured on a Nikon AR1 HD25 confocal microscope using NIS-Elements C Imaging software v5.20.01, and maximum intensity projections were generated in ImageJ Fiji. Mushroom body morphological defects were called as α-lobe thinning or missing and β-lobe fusion or missing for *control, Nab2*^*ex3*^, and control and experimental PCP alleles (e.g., *Vang*^*stbm-6*^*/+, appl*^*d*^*/+*, and *dsh*^*1*^*/+* paired with *control* or *Nab2*^*ex3*^). Statistical analyses for MB phenotypes and plotting performed using GraphPad Prism8^™^. Significance determined using student’s t-test or ANOVA as indicated in figure legends. Error bars representing standard deviation. Significance scores indicated are \**p*≥0.05, \*\**p*≥0.01, and \*\*\**p*≥0.001. Vang-eGFP fluorescence intensity was quantified using two isolated regions of interest (ROI). One ROI located at right hemisphere dorsal cortical surface above the α-lobe (referred to as cortical surface ROI) and a second ROI located at left hemisphere central complex neuropil approximately near the β-lobe and ellipsoid body (referred to as central neuropil ROI). The fluorescence intensity of each ROI was measured in *control* (n=9) and *Nab2*^*ex3*^ (n=9) brains. Significance determined using student’s t-test; significances scores indicated by * = *p*<0.05.

### *Drosophila* neuron live imaging confocal microscopy, neuronal reconstruction, data analyses, and statistical analysis

Live imaging of class IV dorsal dendritic arbor C (ddaC) neurons was performed essentially as described as described in (Iyer *et al*. 2013; Clark *et al*. 2018). Briefly, 3rd instar *ppk-Gal4,mCD8::GFP* labelled larvae were mounted in 1:5 (v/v) diethyl ether:halocarbon oil under an imaging bridge of two 22×22mm glass coverslips topped with a 22×50mm glass coverslip. ddaC images were captured on an Olympus FV 1000 BX61WI upright microscope using Olympus Fluoview software v4.2. Maximum intensity projections were generated with ImageJ Fiji. Neuronal reconstruction was performed with the TREES toolbox (Theisen *et al*. 1994). MathWorks Matlab R2010a v7.10.0.499 (Natick, MA) was used to process 2D stacks with local brightness thresholding, skeletonization, and sparsening to leave carrier points (Cuntz *et al*. 2010). Dendritic roots were defined at the soma and used to create synthetic dendritic arbors. Reconstruction parameters were equivalent across neurons. Various morphological metrics were obtained using the TREES toolbox including: Sholl analysis, total cable length, maximum path length, number of branch points, mean path/Euclidean distance, maximum branch order, mean branch order, mean branch angle, mean path length, mean branch order, field height/width, center of mass x, and center of mass y. These metrics were extracted in batch processing using in-house custom scripts and exported into RStudio v1.1.453 (Vienna, Austria), where quantification was visualized using other in-house custom scripts. Statistical analyses for ddaC phenotypes and plotting were performed using RStudio and Matlab. Balloon plots showing phenotypic data generated using either ddaC measurements generated in Matlab or MB defect counts. Balloon plots generated using RStudio v1.1.453 ggpubr v0.2 (Alboukadel 2018; R-TEAM 2018).

### Global proteomics

MS/MS-LC data was previously described in (Corgiat *et al*. 2021). Briefly, ten biological replicates of 24 hr apf control (*w*^*1118*^) or *Nab2*^*ex3*^ pupal brains (60 brains per replicate) were lysed in urea buffer (8 M urea, 100 mM NaHPO4, pH 8.5) with HALT protease and phosphatase inhibitor (Pierce/Thermo Scientific) and processed at the Emory Proteomics Core. Separate samples were prepared for male and female brains. Label-free quantification analysis was adapted from a previously published procedure (Seyfried *et al*. 2017). Data was analyzed using MaxQuant v1.5.2.8 with Thermo Foundation 2.0 for RAW file reading capability. Spectra were searched using the search engine Andromeda and integrated into MaxQuant against the *Drosophila melanogaster* Uniprot database (43,836 target sequences). Analyses presented here used RStudio v1.1.453 (R-TEAM 2018), custom in-house scripts, and the following packages: ggpubr v0.2 (Alboukadel 2018), cluster v2.1.0 (Maechler *et al*. 2016), and GOplot v1.0.2 (Walter *et al*. 2015), to examine ‘planar cell polarity’ annotated proteins. Gene ontology analyses were performed using FlyEnrichr (FlyEnrichr:amp.pharm.mssm.edu/FlyEnrichr/) (Chen *et al*. 2013; Kuleshov *et al*. 2016; Kuleshov *et al*. 2019), a *Drosophila* specific gene ontology enrichment analysis package.

### Mouse strain, animal care, and histologic analysis of inner ear tissues

The *Zc3h14*^*Δex13/Δex13*^ mouse line has been (referred to as *Zc3h14*^*Δ13/Δ13*^ or *Δ13/Δ13*) described previously (Rha *et al*. 2017). Generations F4-F8 of *Zc3h14*^*Δex13/Δex13*^ backcrosses were used with *Zc3h14*^*+/+*^ controls. All procedures involving mice were done in accordance with HIH guidelines and approved by the Emory University Institutional Animal Care and Use Committee. Cochlea dissection, sectioning, and immunostaining from E14.5 animals has been described previously in (Radde-Gallwitz *et al*. 2004).Whole-mounts of organs of Corti were stained with FITC-conjugated phalloidin to label the stereocilia as in (Wang *et al*. 2005; Qian *et al*. 2007) and samples were analyzed and imaged using a Zeiss LSM510. Cochlear morphological defects were called as extra based on a three OHC layers and one ICH layer separated by pillar cell region. Significance determined using student’s t-test; significance scores indicated by \**p*<0.05

## Results

### Nab2 loss alters levels of planar cell polarity pathway proteins in the *Drosophila* brain

Our recent study comparing proteomic changes in *Drosophila* pupal brains lacking Nab2 identified *planar cell polarity* gene ontology (GO) terms as one category of significantly altered factors (Corgiat *et al*. 2021) (**Fig. 1A**). A deeper analysis of this protein dataset detects enrichment of five PCP-related GO terms (*establishment of planar polarity, establishment of epithelial cell planar polarity, establishment of body hair or bristle planar polarity, protein localization involved in planar polarity, regulation of establishment of planar polarity*) (**Fig. 1B**) extracted from 17 PCP-annotated proteins, including the core PCP component Van Gogh (Vang), and five putative PCP effectors: the Tumbleweed GTPase activating protein (GAP) (Sotillos and Campuzano 2000; Jones *et al*. 2010), the neuron-specific PCP modulator Appl (Singh and Mlodzik 2012; Soldano *et al*. 2013; Liu *et al*. 2021), the anchoring protein Akap200 (Jackson and Berg 2002; Weber *et al*. 2012; Bala Tannan *et al*. 2018), the endocytic regulator X11Lβ (Gross *et al*. 2013), and the muscle LIM-domain protein at 84B (Mlp84B) (Weber *et al*. 2012). Together these factors represent 6.4% of the total differentially expressed proteins in *Nab2*^*ex3*^ pupal brains relative to control (346 proteins in total) (see Corgiat *et al*. 2021) (**Table S1**). The Vang protein (decreased 5-fold in *Nab2*^*ex3*^ vs control) and Appl protein (increased 1.5-fold in *Nab2*^*ex3*^ vs control) are particularly notable because alleles of these genes dominantly modify phenotypes produced by *GMR-Gal4* driven Nab2 overexpression in the developing retinal field (Lee *et al*. 2020).

**Figure 1:**
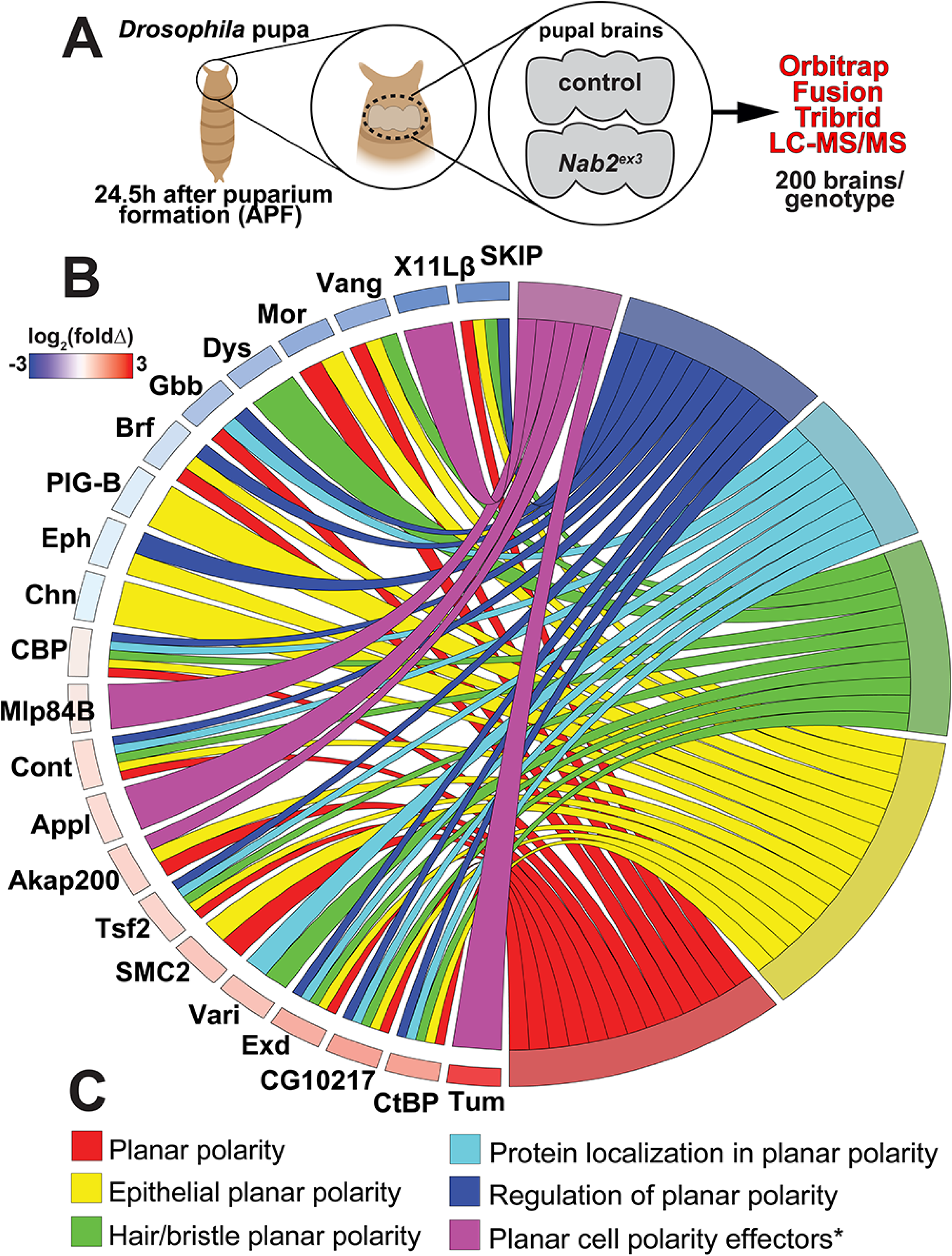
Nab2 loss alters levels of planar cell polarity pathway proteins in the *Drosophila* brain. **(A)** Schematic summary of quantitative proteomic analysis of *Nab2*^*ex3*^ pupal brains dissected from *control* or *Nab2*^*ex3*^ pupa 24.5 hours after puparium formation. Ten samples per genotype, each composed of 20 brains (i.e., 200 *control* brains and 200 *Nab2*^*ex3*^ brains) were processed and analyzed using an Orbitrap Fusion Tribrid Mass Spectrometer and data was quantified using MaxQuant against the *Drosophila melanogaster* Uniprot database. **(B)** Chord plot analysis of protein abundance changes in *Nab2*^*ex3*^ relative to *control* for selected color-coded planar cell polarity ontology terms. Heat map indicates fold-change in abundance of each protein (log_2_(*Nab2*^*ex3*^/*control*)); blue=decreased, red=increased.

### Planar cell polarity components dominantly modify Nab2 axonal phenotypes

To pursue the Nab2-PCP link in the developing CNS, we tested whether axon projection defects in MBs homozygous for the *Nab2*^*ex3*^ null allele (Pak *et al*. 2011) are sensitive to subtle modulation of PCP pathway activity using single copies of loss-of-function alleles of PCP components. Our previous work established genetic interactions between Nab2 and an array of PCP/Wnt alleles in the adult *Drosophila* eye (Lee *et al*. 2020). Here, we focused on three of these factors: the core PCP/Wnt factor Vang, which proteomic data indicate is reduced 5-fold in *Nab2*^*ex3*^ brains (Corgiat *et al*. 2021), the accessory factor Appl (Amyloid precursor protein-like), which is a proposed PCP/Wnt co-receptor and has established links to neurological disease (Singh and Mlodzik 2012; Soldano *et al*. 2013; Liu *et al*. 2021), and the PCP/Wnt cytoplasmic adaptor Dsh, which also genetically interacts with *Nab2* in the wing to control hair polarity (Lee *et al*. 2020). As has been observed in *Nab2*^*ex3*^ adult brains (Kelly *et al*. 2016; Bienkowski *et al*. 2017), *Nab2*^*ex3*^ mutant pupal brain at 48-72h APF (<u>a</u>fter <u>p</u>uparium formation) display highly penetrant defects in structure of the α-lobes (85% thinned or missing) and β-lobes (88% fused or missing) as detected by anti-Fas2 staining (**Fig. 2A-D,Q,R**). Both the *Vang*^*stbm6*^ and *Appl*^*d*^ loss-of-function alleles have no effect on MB structure in an otherwise wildtype background but suppress the frequency of *Nab2*^*ex3*^ α-lobe defects from 85% to 49% in a *Vang*^*stbm6*^*/+* heterozygous background and to 62% in a *Appl*^*d*^*/+* heterozygous background; the frequency of *Nab2*^*ex3*^ β-lobe defects drops from 88% to 33% in *Vang*^*stbm6*^*/+* heterozygous background and to 35% in *Appl*^*d*^*/+* heterozygous background (**Fig. 2E-F,I-J,M-N**). The PCP-specific allele *dsh*^*1*^ (Theisen *et al*. 1994; Gombos *et al*. 2015) lowers *Nab2*^*ex3*^ α-lobe defects from 85% to 63% but has no effect on the frequency or severity of *Nab2*^*ex3*^ β-lobe defects (**Fig. 2Q,R**) (**Fig. S1**). Intriguingly, animals with single copies of *Vang*^*stbm6*^, *Appl*^*d*^, and *dsh*^*1*^ in the *Nab2*^*ex3*^ homozygous background also develop a MB phenotype not observed in any single mutant: a bulbous, Fas2-positive lobe at the point at which the peduncle splits into the five lobes (α,α’,β,β’,γ) (arrowhead in **Fig. 2G,K,O**). The basis of this bulbous phenotype is unclear but may indicate that lowering levels of PCP factors in Kenyon cells that also lack Nab2 leads to a novel axon guidance defect among α/β axons. In sum, these data reveal a pattern of dose-sensitive genetic interactions between endogenous Nab2 and PCP factors that are consistent with Nab2 modulating PCP-mediated control of MB axon projection.

**Figure 2:**
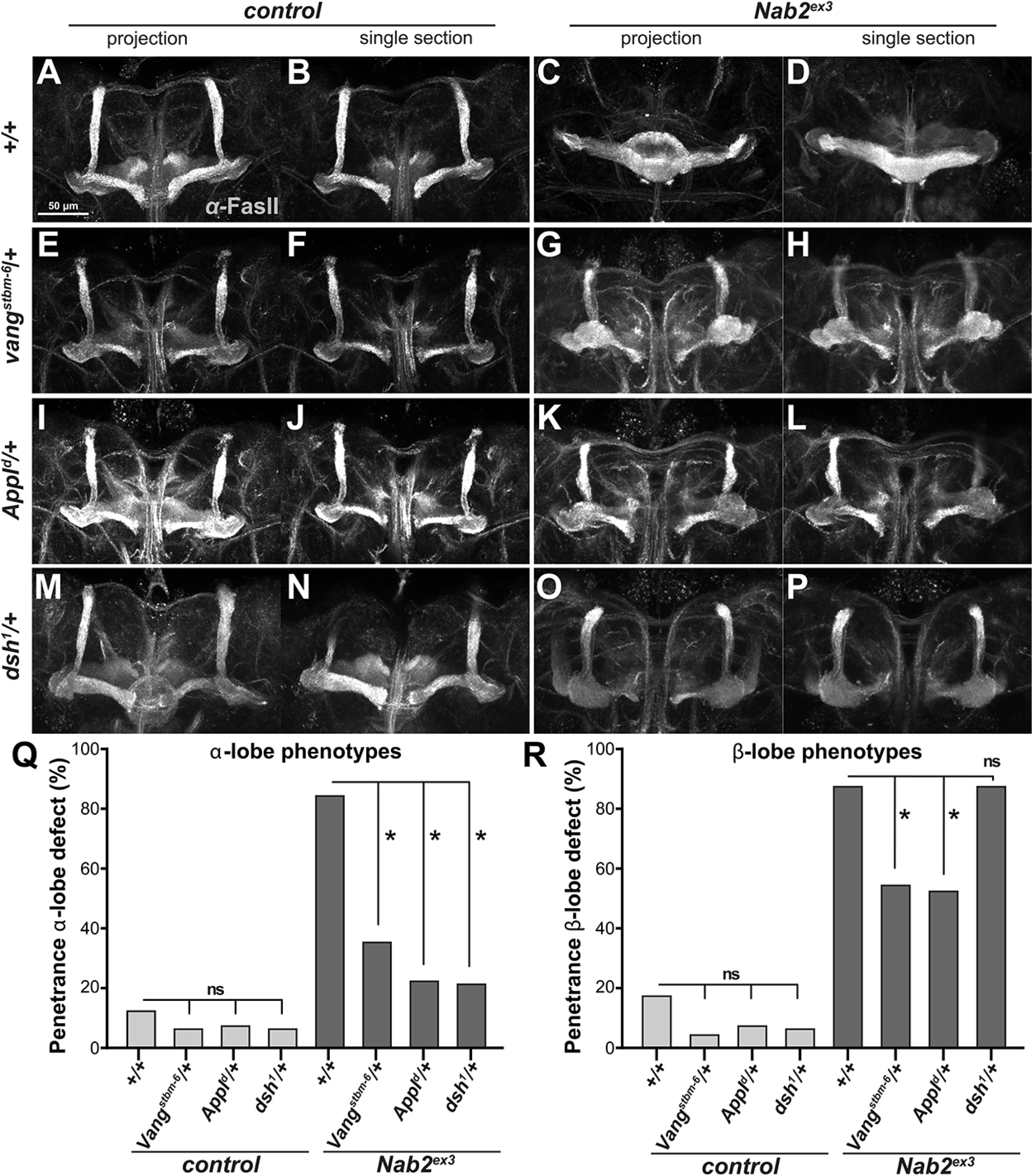
Planar cell polarity components dominantly modify Nab2 axonal phenotypes. Paired maximum intensity Z-stack projections images and single transverse sections of anti-Fasciclin II (FasII) stained 48-72hr pupal brains from **(A-B)** *control* or (**C-D**) *Nab2*^*ex3*^ animals, or each of these genotypes combined with (**E-H**) *Vang*^*stbm6*^*/+*, (**I-L**) *Appl*^*d*^*/+*, or (**M-P**) *dsh*^*1*^*/+*. Frequencies of **(Q)** α- lobe or **(R)** β-lobe structure defects in these genotypes using the scoring system as described in Experimental Procedures. *Nab2*^*ex3*^ brains show high penetrance thinning/loss of α-lobes (85%) and fusion/missing of β-lobe (88%) which are dominantly suppressed by *Vang*^*stbm-6*^ (49% α-lobe and 33% β- lobe defects), *Appl*^*d*^ (62% α-lobe and 35% β-lobe defects). *dsh*^*1*^ selectively suppress *Nab2*^*ex3*^ α-lobe defects to 63%.

### Nab2 is required to restrict dendritic branching and projection

Loss of murine *Zc3h14* causes defects in dendritic spine morphology among hippocampal neurons (Jones *et al*. 2021) prompted us to test whether Nab2-PCP interactions in axons are also conserved in developing dendrites. For this approach, we visualized dendrites of *Drosophila* class IV dorsal dendritic arbor C (ddaC) neurons located in the larval body wall using a *pickpocket (ppk)-Gal4,UAS-GFP* system and quantified branching using Sholl intersection analysis (**Fig. 3F**) (Cuntz *et al*. 2010). In L3 larvae, complete loss of Nab2 leads to increased dendritic branch complexity measured by the number of Scholl intersections relative to control (median of 200 in *ppk>GFP* vs. median of 252 in *Nab2*^*ex3*^; **Fig. 3A-B,G**) which is phenocopied by Nab2 RNAi depletion in ddaC neurons (median of 250 intersections in *ppk>Nab2*^*RNAi*^; **Fig. 3C,G**). Nab2 overexpression in ddaC neurons using the *Nab2*^*EP3716*^ transgene has the inverse effect of decreasing Scholl intersections (median of 179 in *ppk>Nab2*; **Fig. 3E,G**). Significantly, RNAi depletion of the Wnt/PCP receptor *frizzled 2* in ddaC neurons also increases Scholl intersections (median of 216 in *ppk>fz2*^*RNAi*^; **Fig. 3D,G**), confirming prior work that Wnt/PCP signaling is involved in ddaC dendritic development (Misra *et al*. 2016). Significantly, these effects of Nab2 loss on dendritic complexity increase with distance from the cell body (**Fig. 3H**), suggesting that the role of Nab2 in dendritic development becomes more significant with increasing distance from the cell soma.

**Figure 3:**
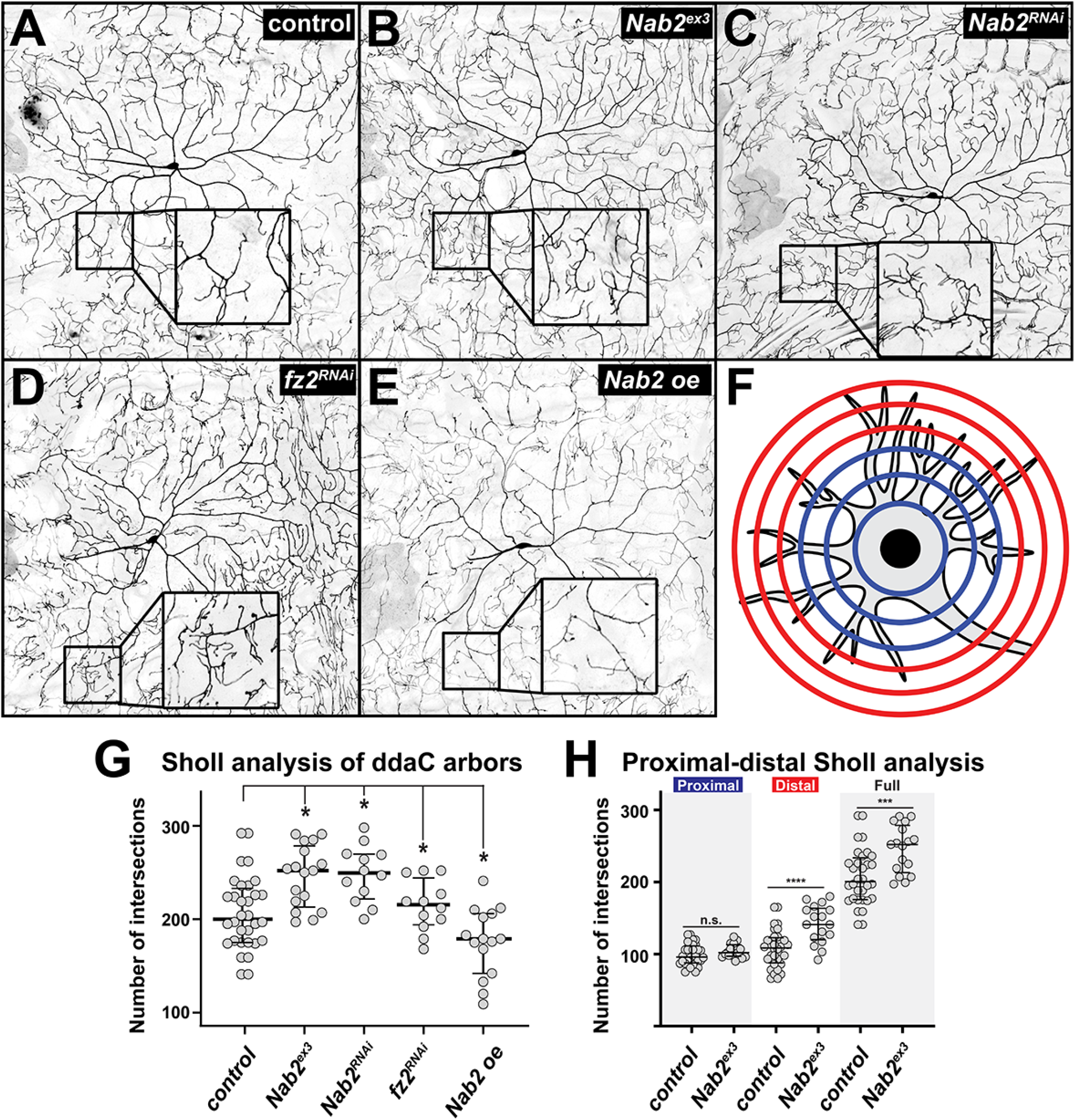
Nab2 is required for proper dendritic development. Inverted intensity images of *Drosophila* class IV dorsal dendritic arbor C (ddaC) neurons from (**A**) *pickpocket (ppk)-Gal4,UAS-GFP*, (**B**) *Nab2*^*ex3*^, (**C**) *ppk-Gal4,UAS-GFP,Nab2*^*RNAi*^, (**D**) *ppk-Gal4,UAS-GFP,fz2*^*RNAi*^, and (**E**) *ppk-Gal4,UAS-GFP,Nab2*^*oe*^ L3 larvae. Inset black boxes show high magnification views of dendritic arbors. (**F)** Diagram depicting the concentric rings used to perform Sholl analysis overlaid on the dendritic arbor of a neuron. The half of the rings proximal to the soma labeled in blue; the half of the rings distal to the soma labeled in red. (**G-H**) Quantification of branching complexity by Sholl analysis of total intersections across dendritic arbor; bars represent median and upper/lower quartile, * *p*<0.05. (**G**) Sholl analysis of full dendritic arbor. Median Sholl intersection values are 200 in *ppk-Gal4,UAS-GFP* (n=32), 252 in *Nab2*^*ex3*^ (n=17), 250 in *ppk-Gal4,UAS-GFP,Nab2*^*RNAi*^ (n=12), 216 in *ppk-Gal4,UAS-fz*^*RNAi*^ (n=12), and 179 in *ppk-Gal4,UAS-Nab2*^*oe*^ (n=15). (**H**) Sholl analysis of proximal and distal dendritic arbors. Median Sholl intersection values for *ppk-Gal4,UAS-GFP* (n=32) are 96 proximal and 108.5 distal, while median Sholl intersection values for *Nab2*^*ex3*^ are 102 proximal and 141 distal.

The data above confirm that Nab2 and the PCP pathway are each required within ddaC neurons to guide the degree of dendritic branching. To further assess whether modulation of PCP pathway activity affects this newly defined *Nab2* dendritic role, we exploited the Matlab TREES toolbox and custom scripts to simultaneously quantify multiple dendritic phenotypes in *Nab2*^*ex3*^ homozygous larvae (**Fig. 4E**) (Cuntz *et al*. 2010). This approach confirmed that *Nab2* loss elevates the total number of branches compared to control (**Fig. 4A,B,D**), but also revealed an extension of overall cable length (**Fig. 4A,B,C**) indicative of increased total projections. A further breakdown of *Nab2*^*ex3*^ branching patterns shows an increase in maximum branch order (# of branch points along a given branch from soma to distal tip) (**Fig. 4D,F**) and coupled decrease in mean branch length (distance between consecutive branches) (**Fig. 4F**). Thus, *Nab2*^*ex3*^ ddaC arbors project and branch significantly more than control across multiple parameters (**Fig. 4F**). Due to the increased branching, *Nab2*^*ex3*^ ddaC arbors exhibit reduced mean path length (−4%), smaller mean branch angles (−9%), and smaller mean branch lengths (−22%) compared to control (**Fig. 4F**). Significantly, these effects of Nab2 loss on branch and length metrics increase with distance from the cell body (**Fig. S2A-B**), which is consistent with a model in which Nab2 restriction of dendrite growth and branching is more significant with increasing distance from the cell soma.

**Figure 4:**
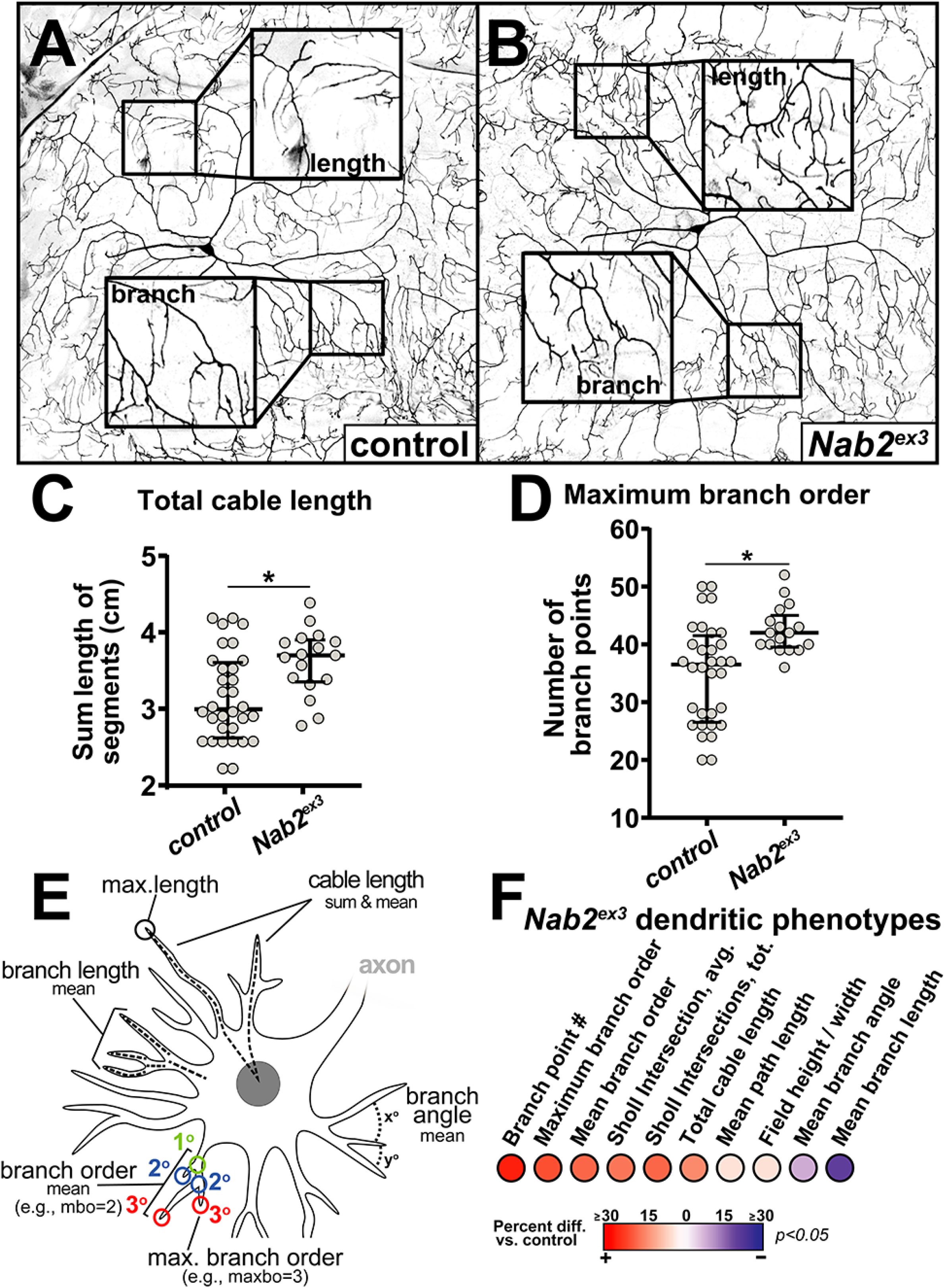
Nab2 restricts dendritic branching and projection. Inverted intensity images of *Drosophila* class IV dorsal dendritic arbor C (ddaC) neurons from (**A**) *control +/+*, (**B**) *Nab2*^*ex3*^ larvae. Inset black boxes show high magnification views of dendritic arbors. Quantification of **(C)** total cable length and **(D)** maximum branch order for *control* (n=32) and *Nab2*^*ex3*^ (n=17); bars represent median and upper/lower quartile, * *p*<0.05. **(E)** Schematic depicting measured dendritic parameters using Matlab TREES toolbox and custom scripts. **(F)** Balloon plot depicting ten measurements of the *Nab2*^*ex3*^ dendritic arbor. Heat map shows change percent changes in *Nab2*^*ex3*^ vs *control*.

### Planar cell polarity components dominantly modify Nab2 dendritic phenotypes

Having established that Nab2 loss elicits a spectrum of ddaC branching and projection defects, we proceeded to test whether genetic modulation of PCP activity could affect one or more of these parameters. While *Nab2*^*ex3*^ homozygotes show an increase in arborization compared to controls (**Fig. 5A-B**), single copies of the *Vang*^*stbm6*^ and *Appl*^*d*^ alleles (i.e., as heterozygotes) each have no significant effects on ddaC arbors in an otherwise wildtype background. In contrast, *dsh*^*1*^ heterozygosity results in increased branch points, Sholl intersections, and total cable length compared to controls. When placed into the *Nab2*^*ex3*^ background, single copies of *Vang*^*stbm6*^ and *Appl*^*d*^ alleles dominantly modify *Nab2*^*ex3*^ phenotypes in opposite directions: *Vang*^*stbm6*^ enhances the severity of *Nab2*^*ex3*^ ddaC branching and length phenotypes while *Appl*^*d*^ suppresses many of the same phenotypes (e.g., total cable length and maximum branch order; **Fig. 5I-K**). The *dsh*^*1*^ allele enhances *Nab2*^*ex3*^ phenotypes (**Fig. 5I-K**), although the presence of ddaC defects in *dsh*^*1*^ heterozygotes suggests that this could be an additive effect. Intriguingly, use of Matlab TREES to assess branching defects as a function of distance from the cell body indicates that complexity changes induced by the *Vang*^*stbm6*^ allele are primarily in *Nab2*^*ex3*^ proximal arbors, while those associated with *Appl*^*d*^ are primarily in distal areas of *Nab2*^*ex3*^ arbors (**Fig. S2B**). Collectively, these genetic and quantitative data argue that Nab2 and PCP components are each individually required for control of ddaC arbors, and that loss of *Nab2* sensitizes ddaC development to the dosage of the core PCP component Vang and the accessory component Appl.

**Figure 5:**
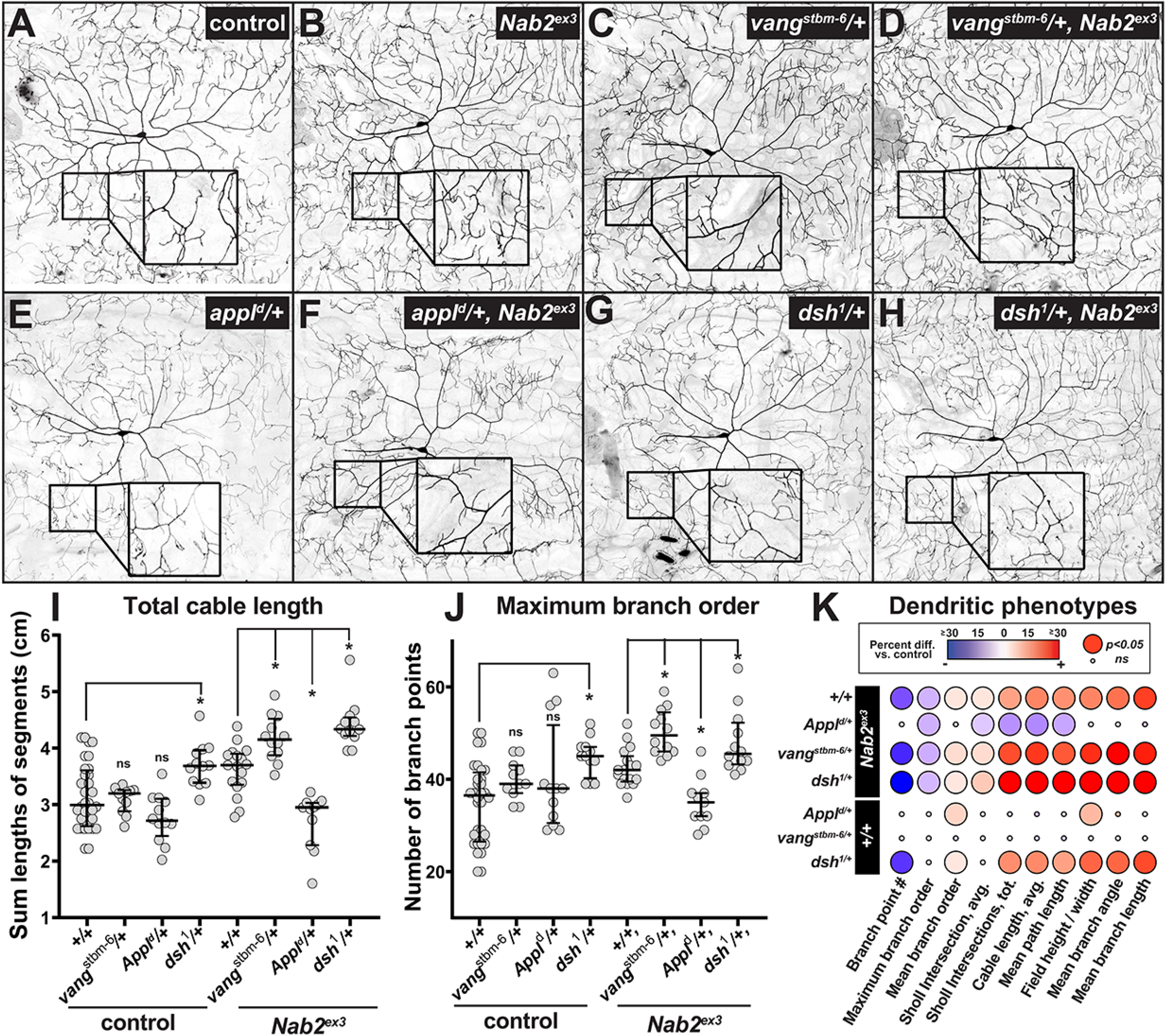
Planar cell polarity components dominantly modify Nab2 dendritic phenotypes. Inverted intensity images of *Drosophila* class IV dorsal dendritic arbor C (ddaC) neurons from (**A**) *control +/+* or **(B)** *Nab2*^*ex3*^ larvae alone, or in combination with **(C-D)** *Vang*^*stbm6*^*/+*, **(E-F)** *Appl*^*d*^*/+*, **(G-H)** *dsh*^*1*^*/+*. Inset black boxes show high magnification views of dendritic arbors. **(I-J)** Quantification of **(I)** total cable length and **(J)** maximum branch order in the indicated genotypes; errors bars represent median and upper/lower quartile, \**p*<0.05. **(K)** Balloon plot analysis of 10 arbor parameters in the indicated genotypes. Heat map shows change percent changes in *Nab2*^*ex3*^ vs *control*. Significance depicted by balloon size (large balloon = p<0.05, small balloon = ns).

### Nab2 is required for proper Vang localization in the central complex of the brain

The pattern of genetic interactions between *Nab2* and *Vang* alleles across the axon-dendrite axis parallel the tandem mass-spectrometry (MS-MS) detection of reduced levels of Vang protein in *Nab2*^*ex3*^ fly brains (**Fig. 1** and see also (Corgiat *et al*. 2021)) and altered levels of Vangl2 protein in *Zc3h14* knockout murine brains (Rha *et al*. 2017). Given these data, we analyzed Vang protein distribution in control and *Nab2*^*ex3*^ brains using a *P[acman]* genomic fragment containing the complete *Vang* locus with an *eGFP* inserted at the C-terminus of the coding sequence and retaining endogenous 5’ and 3’UTRs (*Vang*^*eGFP*.C^) (Strutt *et al*. 2016). This *Vang*^*eGFP*.C^ transgene rescues *Vang* loss-of-function phenotypes and thus provides a reliable readout of Vang expression patterns. Developmentally timed pupal brains were analyzed for Vang-eGFP (anti-GFP) and Bruchpilot (Brp), a presynaptic active zone protein highly enriched in the neuropil (Wagh *et al*. 2006). In control brains, Vang-eGFP fluorescence is distributed in cell bodies at the brain cortical surface as well throughout the Brp-positive central complex brain neuropil, which represents Vang in axons, dendrites, and glial processes (**Fig. 6A-B, D-E**). In contrast, Vang-eGFP is absent in Brp-positive neuropil regions of *Nab2*^*ex3*^ brains (**Fig. 6D,F**) but is present in cortical surface cell bodies and other areas of the brain, including the intersection of the lateral anterior optic tubercle and medulla layer (Neriec and Desplan 2016; Krzeptowski *et al*. 2018; Tai *et al*. 2021) (**Fig. S4B**,**E**) and ventral cortical surface adjacent to the antennal lobes (Wolff *et al*. 2015; Wolff and Rubin 2018) (**Fig. S4D-G**). Quantification of Vang-eGFP signal intensity in Brp-positive central neuropil (‘n’ box region in **Fig. 6B,E**) and a region of the dorsal cortical surface (‘c’ box in **Fig. 6B,E**) reveals substantial loss of neuropil-localized Vang-eGFP in *Nab2*^*ex3*^ brains relative to controls, with no significant effect on the level of cortical Vang-eGFP in cell bodies (**Fig 6G-H**). Given that Brp-positive neuropil regions are enriched in axons, dendrites, and glial processes, these data indicate that Nab2 is required for Vang-eGFP protein to accumulate in distal neuronal and glial processes, and that the overall drop in Vang protein levels detected in MS-MS analysis of *Nab2*^*ex3*^ brains (**Fig. 1B**) is accompanied by a change in steady-state Vang-eGFP localization that may deprive distal axon-dendritic compartments of factors required for normal PCP signaling.

**Figure 6:**
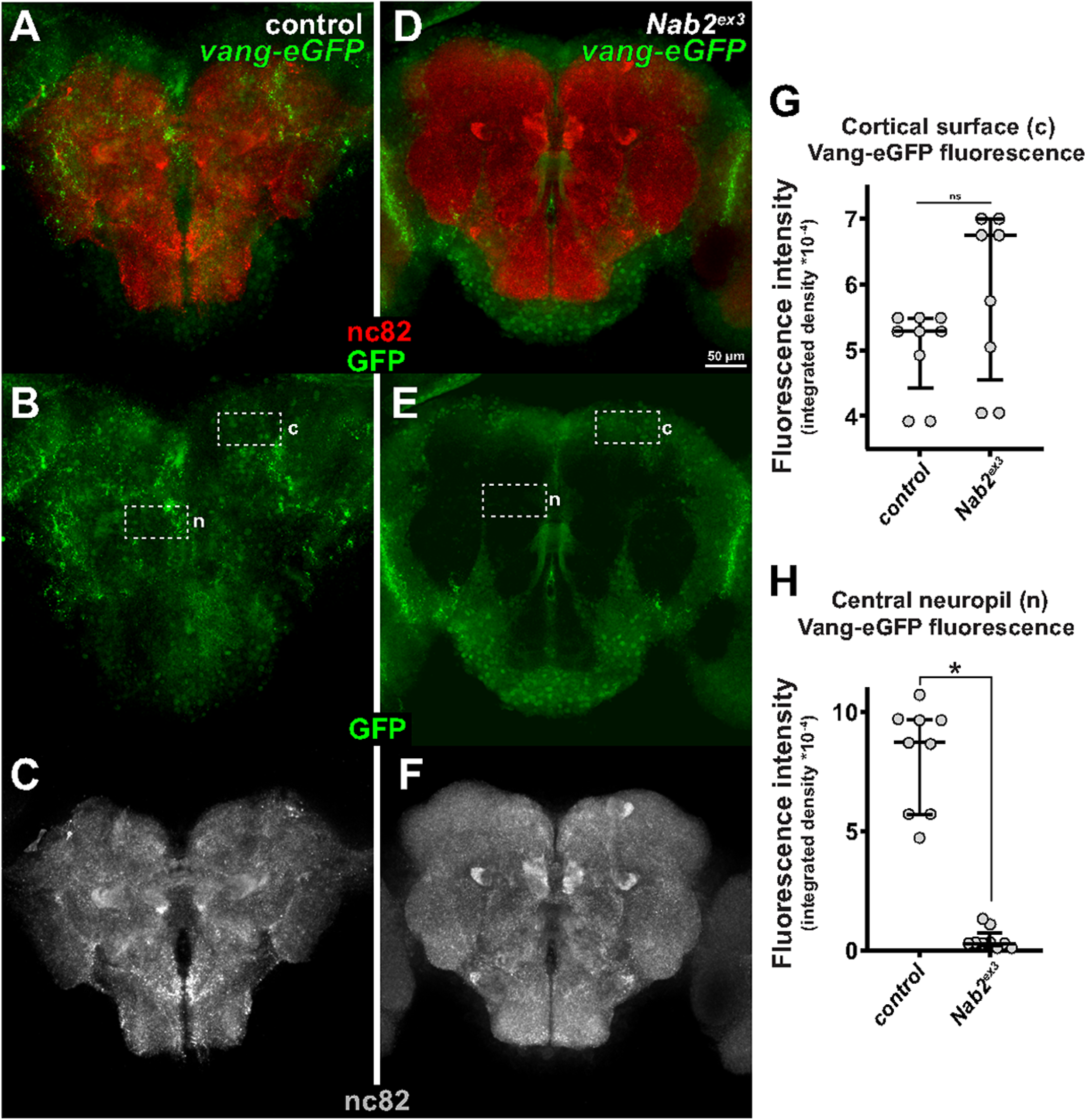
Nab2 is required for proper Vang localization in the central complex of the brain. Visualization of brains from 48-72hr *vang*^*EGFP*.C^ (**A-C**) and *Nab2*^*ex3*^*;vang*^*EGFP*.C^ (**D-F**) pupae co-stained with anti-GFP (green) and the nc82 mAb (red) to detect Vang-eGFP and Brp, which marks presynaptic actives zones. Dashed boxes indicate regions used for quantifying fluorescence in **c** (cortical surface) and **n** (central neuropil) regions. (**G-H**) Quantification of Vang-eGFP fluorescence intensity in the (**G**) cortical surface and (**H**) central neuropil regions of *vang*^*EGFP*.C^ (n=9) and *Nab2*^*ex3*^*;vang*^*EGFP*.C^ (n=9) pupae. Bars represent median and upper/lower quartile, * *p*<0.05.

### *Zc3h14* deficient mice show PCP-like defects in the cochlea

Many phenotypes are conserved from *Nab2*^*ex3*^ flies to *Zc3h14*^*Δ13/Δ13*^ mice including defects in working memory (Kelly *et al*. 2016; Bienkowski *et al*. 2017; Rha *et al*. 2017), a subset of proteomic changes in the brain (Rha *et al*. 2017; Corgiat *et al*. 2021), and defects in dendritic morphology (this study and Jones *et al*. 2021). Nab2 has strong genetic interactions with PCP components, as shown here, but also has PCP-like defects in orientation of the fly wing hair bristles, a classic PCP phenotype (Adler 2012; Olofsson and Axelrod 2014; Lee *et al*. 2020). Given that mammalian ZC3H14 can rescue a variety of phenotypes when expressed in the neurons of *Nab2* mutant *Drosophila* (Pak *et al*. 2011; Kelly *et al*. 2014; Bienkowski *et al*. 2017), we assessed *Zc3h14*^*Δ13/Δ13*^ mice for evidence of PCP defects, with a focus on elements of the sensory nervous system. Development of the organ of Corti within the cochlea is well established as a PCP-regulated process in the mouse (Jones and Chen 2007; Chacon-Heszele and Chen 2009; Rida and Chen 2009). The organ of Corti is formed by sensory cells, known as hair cells, that are patterned in one row of inner cells, and three rows of outer cells (Chacon-Heszele and Chen 2009). Mutations in murine PCP genes result in altered orientation and patterning of these hair cells, due in part to the requirement for PCP in the process of convergent extension. To test whether a Nab2-PCP functional link is conserved in the mouse cochlea, we analyzed *Zc3h14*^*Δ13/Δ13*^ mutant cochlea for PCP-like phenotypes. Phalloidin staining the organ of Corti from E14.5 *Zc3h14*^*Δ13/Δ13*^ embryos revealed additional rows of outer hair cells (OHCs) in both the basal and middle regions compared to control (**Fig. 7A**). There are occasional inner hair cells (IHCs) patterning defects in the middle region (**Fig. 7A**). Quantification of extra cells per cochlea confirmed significant OHC patterning defects (**Fig. 7B**) in *Zc3h14*^*Δ13/Δ13*^ mice with no significant defects in IHC patterning (**Fig. 7C**). These data suggest that PCP-like phenotypes are shared from *Nab2*^*ex3*^ flies to *Zc3h14*^*Δ13/Δ13*^ mice and that Nab2 interactions with PCP components may be conserved in ZC3H14.

**Figure 7:**
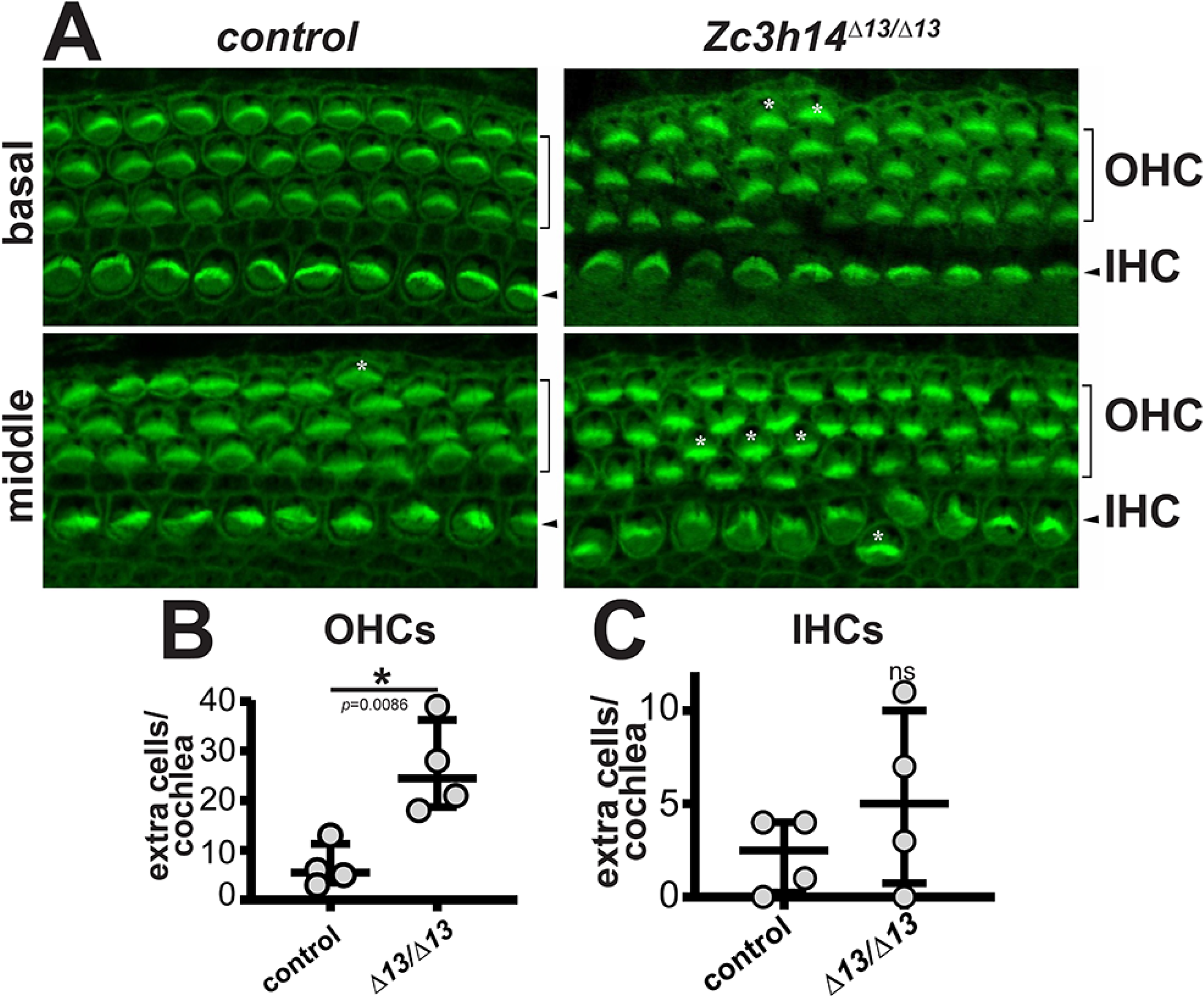
*Zc3h14*^*Δ13/Δ13*^ mice have PCP-like cochlear defects. (**A**) The cochlea from *control* and *Zc3h14*^*Δ13/Δ13*^ E14.5 embryos showing basal and middle regions. Stereocilia are visualized by phalloidin staining. Brackets indicate outer hair cells (OHC) and arrowheads indicate inner hair cells (IHC). Staining reveals normal orientation and hair cell layers for *control* but extra OHC and some orientation defects around the pillar cell region for *Zc3h14*^*Δ13/Δ13*^. (**B-C**) Quantification of extra cells per cochlea in the (**B**) OHC and (**C**) IHC; bars represent median and upper/lower quartile, * indicates p<0.05. *control n*=4, *Zc3h14*^*Δ13/Δ13*^ *n*=4.

## Discussion

Here, we uncover a role for *Drosophila* Nab2, an evolutionarily conserved RBP with links to human inherited intellectual disability, in control of dendrite branching and projection among ddaC body wall sensory neurons via a mechanism that is linked to the Nab2 role in axon projection and branching via shared dependence on the PCP pathway. Loss of Nab2 increases dendrite branching and projection while overexpression of Nab2 has the opposite effect of restricting dendrite growth. Using proteomic data collected from *Nab2* null developing fly brains (Corgiat *et al*. 2021), we uncover an enrichment for planar cell polarity factors among proteins whose steady-state levels are affected by Nab2 loss, and define a pattern of genetic interactions that are consistent with Nab2 regulating projection and branching of ddaC dendrites and MB axons by a common PCP-linked mechanism. Cell type-specific RNAi indicates that Nab2 acts cell autonomously to guide PCP-dependent axon and dendrite growth, implying a direct link between Nab2 and one or more PCP components within ddaC neurons and MB Kenyon cells. Based on reduction in levels of the PCP component Vang detected in our proteomic analysis, we analyze the levels and distribution of a fluorescently tagged Vang protein (Vang-eGFP) in adult brains (Strutt *et al*. 2016). The overall drop in Vang-eGFP levels detected by proteomics is also evident in optical sections of whole brains and is unexpectedly accompanied by selective loss of Vang protein in axon/dendrite-enriched neuropil regions relative to brain regions containing nuclei and cell bodies. Analysis of a *Zc3h14* mutant mouse (Rha *et al*. 2017) reveals PCP phenotypes within the sensory nervous system, suggesting that functional links between Nab2/ZC3H14 and the PCP pathway may be evolutionarily conserved. Collectively, these data demonstrate that Nab2 is required to regulate axonal and dendritic growth by a PCP-linked mechanism and identify the Nab2 RBP as required for the steady-state accumulation of Vang protein in distal neuronal compartments.

RBPs shape axon and dendrite architecture by modulating steps in post-transcriptional regulation of key neuronal mRNAs, including their export, trafficking, stability, and translation (Ravanidis *et al*. 2018; Schieweck *et al*. 2021). Of note, the analysis presented here shows that effects of Nab2 on dendritic morphology are exaggerated in regions of neurons more distal from the soma as compared to more proximal regions (**Fig. 3H; Fig. S2A-B**). This enhanced effect of Nab2 loss on distal branching of ddaC arbors implies that Nab2 controls expression of an mRNA (or mRNAs) encoding a factor that guides branching and projection of more distal dendrites. While neuronal Nab2 protein is primarily nuclear (Pak *et al*. 2011), the protein is also detected in cytoplasmic messenger ribonucleoprotein (mRNP) granules and has a proposed role in translational repression (Bienkowski *et al*. 2017), suggesting that cytoplasmic Nab2 could inhibit translation of mRNAs that traffic to distal dendrites and promote branching and projection. Core PCP proteins localize to membranes at distal tips of some *Drosophila* neuronal growth cones (e.g., Reynaud *et al*. 2015; Misra *et al*. 2016) and multiple *Drosophila* Wnt/PCP factors act autonomously in ddaC cells to control dendritic growth (e.g., *fz2* in this study and see (Matsubara *et al*. 2011)). Considering these observations, Nab2 might inhibit translation of one or more PCP mRNAs, perhaps as it is trafficked for subsequent expression at distal tips of axons and dendrites. The exclusion Vang-eGFP from the axon/dendrite enriched brain neuropil (**Figs. 6** and **S4**) could then be a consequence of failed mRNA transport, followed by precocious translation (and perhaps turnover) in the soma, or it be indicative of a Nab2 role in promoting *Vang* mRNA translation in distal compartments. In sum, these data provide first evidence that *Drosophila* Nab2 may aid in localizing neurodevelopmental factors into distal dendrites, and that this may be coupled with a role in regulating mRNA trafficking and/or translation.

Within brain neurons, Nab2 loss depletes Vang-eGFP from neuropil, which in enriched in axons, dendrites, and glial processes and depleted of soma/nuclei (**Figs. 6** and **S4**). One parsimonious model to explain this observation is that the *Vang-eGFP* mRNA is regulated by Nab2 and that Vang-eGFP depletion from *Nab2*^*ex3*^ brain neuropil is thus due to a defect in *Vang-eGFP* mRNA localization and/or local translation. In this model, Nab2 could either bind directly to the *Vang* mRNA or indirectly regulate *Vang* mRNA via an intermediary factor. As the *Vang*^*eGFP*.*C*^ allele retains the single *Vang* intron and intact 5’ and 3’ UTRs (Strutt *et al*. 2016), post-transcriptional regulation of the *Vang* mRNA by Nab2 should be mirrored by effects on endogenous Vang protein, which indeed drops in abundance in *Nab2* null brains. Intriguingly, Vang protein is expressed in core axons of the α and β MB lobes far from their originating Kenyon cell bodies (Shimizu *et al*. 2011) and is required to pattern these distal axons, as shown by the disruptive effect of *vang* loss on α/β lobe structure (Shimizu *et al*. 2011; Ng 2012). Thus, the interactions between *Nab2* and *Vang* alleles in MB axons may also reflect a specific role for both factors in controlling projection and branching of distal neuronal processes that mirrors their relationship in ddaC dendrites.

The genetic interactions between *Nab2* and alleles of PCP components in MB axons imply a degree of context-dependence to the Nab2-PCP interaction between ddaCs and MBs, and even between the two distinct axon compartments represented by the MB α- and β-lobes. While *Vang*^*sbm6*^ heterozygosity enhances *Nab2*^*ex3*^ ddaC defects, this allele selectively suppresses *Nab2*^*ex3*^ MB α-lobe defects but not β-lobe defects. Given the broad drop in Vang-eGFP levels observed in *Nab2*^*ex3*^ brains (**Fig. 6**), and the requirement for *Vang* in Kenyon cells (KCs) for normal development of both the α and β-lobes (Shimizu *et al*. 2011; Ng 2012), the α-lobe-specific *Nab2-Vang* genetic interaction could be regarded as unexpected. However, very similar α-lobe-specific genetic interactions occur between *Nab2* and alleles of two other RBP genes, *fmr1* and *Atx2* (Bienkowski *et al*. 2017; ROUNDS 2021), implying that Nab2 has distinct interacting pathways in these two different axonal sub-compartments. As noted, suppression of *Nab2*^*ex3*^ MB defects by *Vang*^*stbm6*^ is the inverse of how this same allele affects *Nab2*^*ex3*^ ddaC phenotypes. The relationship could arise if PCP signals are exchanged between MB axons and the surrounding neuro-substrate, which could invert a relationship between Nab2 in Kenyon cells and Wnt/PCP signals emanating from surrounding cells. For example, the receptor *derailed* is expressed in the dorsomedial lineage neuropil and binds Wnt5 for presentation to repulsive *derailed2* receptors on α-lobe axons, thus non-autonomously guiding α-lobe projection (Reynaud *et al*. 2015). In addition, the projection paths of individual *vang*^*stbm6*^ mutant α and β-axon tracts can be rescued by adjacent cells with normal Vang level, indicating that Wnt/PCP control of α and β-axon branching is not strictly cell-autonomous (Shimizu *et al*. 2011; Ng 2012). These complex signaling relationships within MB axons, and potential extra-cellular Wnt/PCP guidance cues emanating from surrounding dorsomedial cells, are both potential explanations for context-specific genetic interactions between *Nab2* and *Vang* in ddaCs and Kenyon cells. In contrast to *Vang* alleles, partial loss of Appl (*Appl*^*d*^) consistently suppresses both *Nab2*^*ex3*^ dendritic and axonal phenotypes (**Fig. S3**) which parallels the increase in Appl protein detected in brain proteomics in *Nab2* mutant brains (**Fig. 1B, Table S1**). Appl acts as a downstream neuronal-specific effector of the PCP pathway (Soldano *et al*. 2013; Liu *et al*. 2021) and elevated Appl protein in response to Nab2 loss could be an indirect consequence of altered core PCP pathway activity or evidence of direct regulation of the *Appl* transcript. These differing interactions illustrate the complexity of RBP function across a neuron with specific interactions affecting sub-cellular compartments in different ways.

An additional question that arises from analysis of Nab2-PCP interactions in the MBs is: *why Nab2 mutant α-lobe defects are rescued by Vang, Appl, and dsh alleles to a greater degree than are β-lobe defects?* As noted above, alleles of the Nab2-interacting factors *fmr1* and *Atx2* also specifically suppress *Nab2*^*ex3*^ α-axon defects but not β-axon defects (Bienkowski *et al*. 2017), implying that these gene may define a Nab2-dependent α-lobe Nab2-Fmr1-Atx2 regulatory network that also includes PCP mRNAs. Indeed, *fmr1* also shares some functional features with *Nab2* in MBs and ddaCs: Fmr1 controls both α- and β-lobe development (Michel *et al*. 2004) and limits ddaC dendrite development, in part through an interaction the mRNA encoding the PCP-effector Rac1 (Fanto *et al*. 2000; Lee *et al*. 2003). Significantly, the normal development of α and β-axons has been proposed to rely on a lobe-specific PCP mechanism involving the formin DAAM (Dsh associated activator of morphogenesis) interacting with Wg/Wnt receptor Frizzled (Fz) in the α-lobes and Vang in the β-lobes (Gombos *et al*. 2015). A similar type of mechanism could occur for the Nab2-PCP interaction, with Nab2 either regulating different PCP components in α vs. β lobes or regulating components that themselves have lobe-specific roles e.g., DAAM or the Derailed-Wnt5 receptor ligand pair (Reynaud *et al*. 2015). In sum, it seems likely that future studies will identify other mechanisms and pathways through which Nab2 regulates axon-dendrite development in opposition to or cooperation with the Wnt/PCP pathway, including for example mechanisms involving the Fmr1 and Atx2 RBPs interacting with Nab2 to regulate expression of co-bound RNAs.

We extended the data from *Drosophila* to mouse by taking advantage of a mouse model lacking functional ZC3H14/Nab2 protein (Rha *et al*. 2017). This analysis reveals that *zc3h14* mutant mice show phenotypes in orientation of the hair cell stereociliary bundles within the cochlea that are similar to multiple PCP mutants, including *Vangl2* (Montcouquiol *et al*. 2003). Future studies could employ mouse models to explore whether genetic interactions identified in Drosophila extend to mammals, but this conserved PCP phenotype argues for a conserved link between ZC3H14/Nab2 and the PCP pathway.

In aggregate, these data reveal Nab2 interactions with the PCP pathway and provide the first evidence that Nab2 is required for dendritic development. These interactions between Nab2 and PCP factors in control of dendritic complexity and MB axon projection appear to be cell-autonomous and, at least in ddaC arbors, more dramatic in distal projections. Changes in expression level and localization of Vang protein in fly brains lacking Nab2 highlight the *Vang* mRNA as a potential target of post-transcriptional control by Nab2 both in axons and dendrites. Given that loss of the Nab2 ortholog in mice, *Zc3h14*, also alters levels of the Vangl2 PCP protein in the adult hippocampus, and that mutations in PCP genes including *Vangl2* are linked to intellectual disabilities, severe neural tube closure defects, and microencephaly in humans (e.g.,Wang *et al*. 2019) dysregulation of the levels and localization of

PCP components in neurons is one potential mechanism to explain axonal and dendritic phenotypes in *Zc3h14* mutant mice (Jones *et al*. 2021) and cognitive defects in human patients lacking ZC3H14 (Pak *et al*. 2011).

## Acknowledgements

We thank Dr. Dan Cox, GA State Neuroscience Institute, for reagents and discussion, and members of the Moberg and Corbett laboratories for helpful discussions. We thank the Emory Proteomics Core for their support and guidance.

## Data availability

Proteomics data have been deposited to the ProteomeXchange Consortium via the PRIDE partner repository with the dataset identifier PXD022984. All remaining data are contained within the article.

## Funding and additional information

Research reported in this publication was also supported in part by the Emory University Integrated Cellular Imaging Microscopy Core of the Emory Neuroscience NINDS Core Facilities grant, 5P30NS055077. Financial support as follows: 5F31NS110312, 5F31HD088043, and 5R01MH107305.

The authors declare no competing financial interests.

